# Signal diversification is associated with corollary discharge evolution in weakly electric fish

**DOI:** 10.1101/2020.04.16.044842

**Authors:** Matasaburo Fukutomi, Bruce A. Carlson

## Abstract

Communication signal diversification is a driving force in the evolution of sensory and motor systems. However, little is known about the evolution of sensorimotor integration. Mormyrid fishes generate stereotyped electric pulses (electric organ discharge [EOD]) for communication and active sensing. The EOD has diversified extensively, especially in duration, which varies across species from 0.1 to over 10 ms. In the electrosensory hindbrain, a corollary discharge that signals the timing of EOD production provides brief, precisely timed inhibition that effectively blocks responses to self-generated EODs. However, corollary discharge inhibition has only been studied in a few species, all with short duration EODs. Here, we asked how corollary discharge inhibition has coevolved with the diversification of EOD duration. We addressed this question by comparing 7 mormyrid species having varied EOD duration. For each individual fish, we measured EOD duration and then measured corollary discharge inhibition by recording evoked potentials from midbrain electrosensory nuclei. We found that delays in the onset of corollary discharge inhibition were strongly correlated with EOD duration as well as delay to the first peak of the EOD. In addition, we showed that electrosensory receptors respond to self-generated EODs with spikes occurring in a narrow time window immediately following the first peak of the EOD. Direct comparison of time courses between the EOD and corollary discharge inhibition revealed that the inhibition overlaps the first peak of the EOD. Our results suggest that internal delays have shifted the timing of corollary discharge inhibition to optimally block responses to self-generated signals.

**SIGNIFICANCE STATEMENT:** Corollary discharges are internal copies of motor commands that are essential for brain function. For example, corollary discharge allows an animal to distinguish self-generated from external stimuli. Despite widespread diversity in behavior and its motor control, we know little about the evolution of corollary discharges. Mormyrid fishes generate stereotyped electric pulses used for communication and active sensing. In the electrosensory pathway that processes communication signals, a corollary discharge inhibits sensory responses to self-generated signals. We found that fish with long duration pulses have delayed corollary discharge inhibition, and that this time-shifted corollary discharge optimally blocks electrosensory responses to the fish’s own signal. Our study provides the first evidence for evolutionary change in sensorimotor integration related to diversification of communication signals.

## INTRODUCTION

Diversification of communication signals is a driving force in animal speciation. Signal evolution has been associated with evolutionary changes to sensory receptors and central sensory circuits (Osorio and Vorobyev, 2008; Carlson et al., 2011; Baker et al., 2015; ter Hofstede et al., 2015; Vélez and Carlson, 2016; Silva and Antunes, 2016; Vélez et al., 2017; Seeholzer et al., 2018), as well as peripheral effectors and central motor circuits (Bass, 1986; Otte, 1992; Podos, 2001; Paul et al., 2015; Ding et al., 2019; Jacob and Hedwig, 2019; Kwong-Brown et al., 2019). Despite this accumulated knowledge of sensory and motor system evolution, we know little about the evolution of sensorimotor interactions between these systems.

Corollary discharges are one of the links by which motor control influences sensory processing to distinguish external from self-generated stimuli (von Holst and Mittelstaedt, 1950; Sperry, 1950; Poulet and Hedwig, 2007; Crapse and Sommer, 2008; Schneider and Mooney, 2018; Straka et al., 2018). For communicating animals, a corollary discharge generally works to filter out an animal’s own signals (reafference), allowing selective sensory processing of signals from other individuals (exafference). Since a corollary discharge needs to selectively cancel reafferent input, its function should evolve to adapt to signal diversification. However, this question has not been addressed to our knowledge.

Here we investigate corollary discharge evolution in mormyrids, African weakly electric fishes. These fish produce electric-pulse signals, termed electric organ discharge (EOD), that are used for electrolocation and communication (Hopkins, 1999; von der Emde, 1999). EOD waveforms are stereotyped but diverse among species and sometimes among individuals within species (Hopkins, 1981; Arnegard et al., 2010; Paul et al., 2015; Gallant and O’Connell, 2020). The EOD is generated from an electric organ (EO) at the base of the tail (Fig. 1A; Bennett, 1971). EOD waveform is determined by the biophysical characteristics of electrocytes in the EO, while EOD timing is determined by central neural commands (Bennett, 1971; Bass, 1986; Carlson, 2002a).

**Figure 1.**
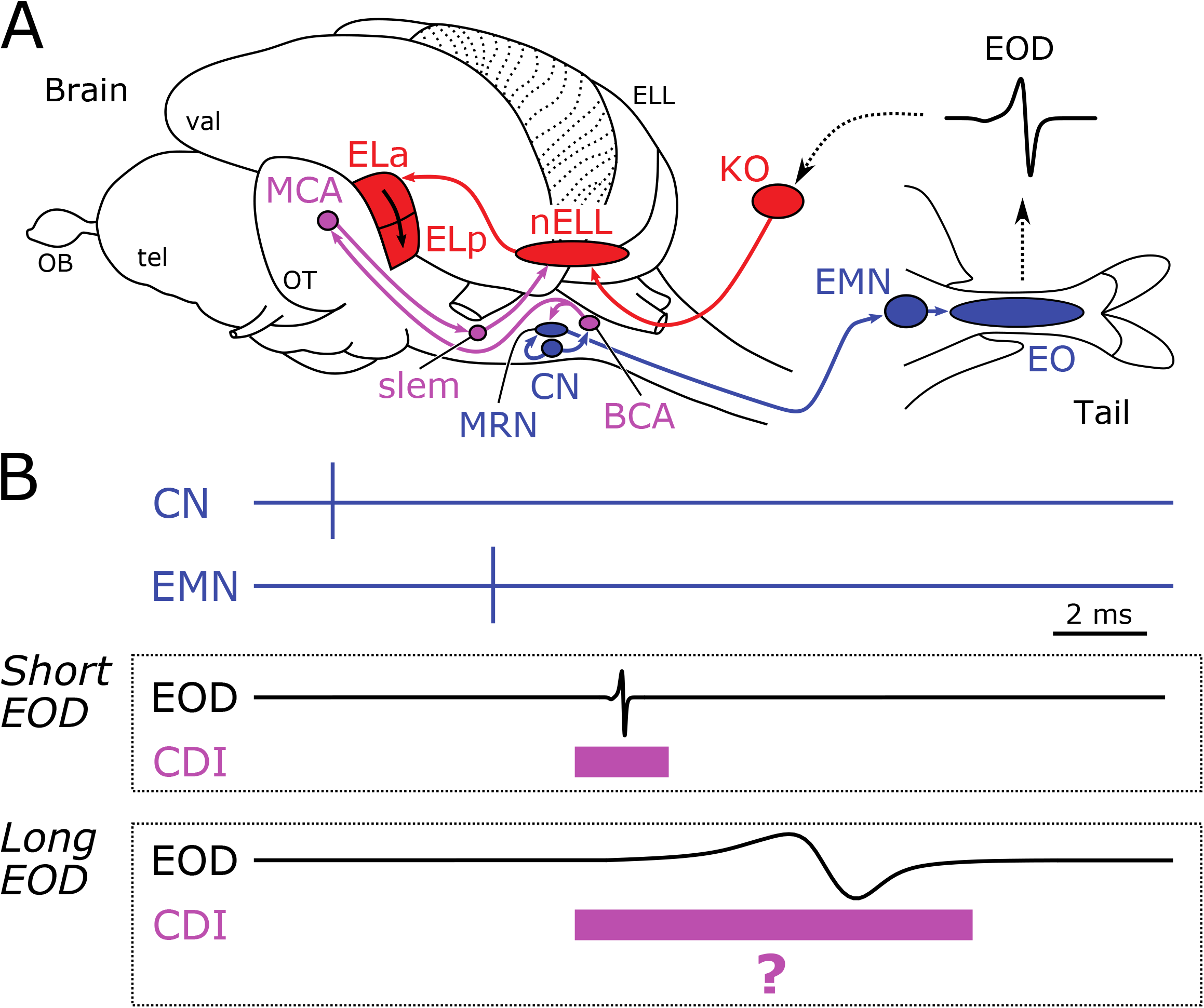
Corollary discharge inhibition and evolution of EOD duration. (A) Diagram showing electromotor (blue), Knollenorgan sensory (red), and corollary discharge (purple) pathways. The command nucleus (CN) drives the EO to generate each EOD via the medullary relay nucleus (MRN) and the spinal electromotor neurons (EMN). Knollenorgan electroreceptors (KO) receive the EOD and send time-locked spikes to the nucleus of the electrosensory lateral line lobe (nELL) via primary afferents. The nELL neurons project their axons to the anterior exterolateral nucleus (ELa), which sends its only output to the adjacent posterior EL (ELp). The CN provides another output to the bulbar command-associated nucleus (BCA), which in turn projects to the MRN and to the mesencephalic command-associated nucleus (MCA). The MCA sends its output to the sublemniscal nucleus that has GABAergic neurons projecting to the nELL. ELL, electrosensory lateral lobe; OB, olfactory bulb; OT, optic tectum; tel, telencephalon; vel, valvula of the cerebellum. (B) Potential hypothesis of corollary discharge inhibition between mormyrids with short EOD and long EOD. The schematic diagram shows spike timings of the CN and EMN as well as EOD. The purple rectangle indicates the potential time window of corollary discharge inhibition (CDI). In the short-EOD species, the corollary discharge covers the KO spike to self-generated EODs so as to inhibit sensory responses in the nELL. Since long-duration EODs can cause KO spikes with different timing, corollary discharge inhibition needs to change its duration and/or timing to block reafferent responses in the nELL.

Electric communication signals are processed by a dedicated sensory pathway (Fig. 1A; Xu-Friedman and Hopkins, 1999; Baker et al., 2013). Sensory receptors called Knollenorgans (KO) respond to outside-positive changes in voltage across the skin, or inward current, with a fixed latency spike (Bell, 1989). Because each KO faces out towards the surrounding water, in response to an external EOD, KOs on one side of the body receive an inward current while KOs on the other side receive an outward current (Hopkins and Bass, 1981). By contrast, in response to a self-generated EOD, KOs on both sides receive the same-direction currents of the waveform consisting of a large outward current followed by a large inward current if the EOD has a head-positive polarity (Fig. 1A). The KO afferent fibers project to the nucleus of the electrosensory lateral line lobe (nELL) in the hindbrain, where corollary discharge inhibition blocks responses to the fish’s own EOD (Fig. 1A; Bell and Grant, 1989). The axons of nELL neurons project to the anterior exterolateral nucleus (ELa) of the midbrain torus semicircularis, which sends its only output to the posterior exterolateral nucleus (ELp) (Fig. 1A; Xu-Friedman and Hopkins, 1999).

Mormyrids are advantageous for studying corollary discharge interactions between motor and sensory systems: the motor command signal (fictive EOD) can be easily recorded from spinal electromotor neurons (EMN) when EOD production is silenced pharmacologically. Previous studies reported that corollary discharge inhibition starts about 2 ms after the onset of command signal and lasts for about 2 ms (Fig. 1B; Amagai, 1998; Lyons-Warren et al., 2013b; Vélez and Carlson, 2016). Those studies used limited species that have short-duration EODs (~0.5 ms), but the mormyrid family has evolved EOD durations ranging from 0.1 to over 10 ms (Hopkins, 1999). The present study uses 7 mormyrid species having EODs that vary in duration from ~0.1 to ~10 ms, and compares corollary discharge inhibition across species to reveal evolutionary change of corollary discharge in relation to signal diversification.

## MATERIALS AND METHODS

All procedures were in accordance with guidelines established by the National Institutes of Health and were approved by the Animal Care and Use Committee at Washington University in St Louis.

### Animals

6 *Brienomyrus brachyistius* (Standard length [SL] = 7.5–11.8 cm), 3 *Brevimyrus niger* (SL = 6.7–9.5 cm), 3 *Campylomormyrus compressirostris* (SL = 11.8–14.0 cm), 6 *Campylomormyrus numenius* (SL = 12.4–14.2 cm), 4 *Campylomormyrus tamandua* (SL = 6.5–9.1 cm), 3 *Gnathonemus petersii* (SL = 10.2–11.6 cm) and 2 *Mormyrus tapirus* (SL = 12.3–12.3 cm) contributed EOD and evoked potential data to this study. We used fish of both sexes in *B. brachyistuis, B. niger, C. numenius* and *C. tamandua*, but only female in *C. compressirostris, G. petersii* and *M. tapirus*. Subsets of these fish (1 *B. brachyistius*, 1 *C. compressirostris* and 1 *C. numenius*) were used for simultaneous recording of the EOD and EOD command generated by EMNs. The fish were housed in water with a conductivity of 175–225 μS/cm, a pH of 6–7, and a temperature of 25–29 °C. The fish were kept on a 12/12-h light/dark cycle and fed live black worms 4 times a week.

### EOD recording and analysis

We recorded 10 EODs from each fish while it was freely swimming. EODs were amplified 10 times, band-pass filtered (1 Hz–50 kHz) (BMA-200, Ardmore), digitized at a rate of 195 kHz (RP2.1, Tucker Davis Technologies), and saved using custom software in Matlab (Mathworks).

EODs generally consist of peak 1 (maximum head-positive peak) and peak 2 (maximum head-negative peak). Some species we recorded from (*B. brachyistius, B. niger, C. tamandua* and *G. petersii*) have EODs with an additional peak 0 (small head negative peak before peak 1). For EODs without a peak 0, EOD onset was determined as the point crossing 2% of peak-1 amplitude. For EODs with a peak 0, EOD onset was determined as the point crossing 20% of peak-0 amplitude. In both cases, EOD offset was determined as the point crossing 2% of peak-2 amplitude. EOD duration was determined as the period between EOD onset and offset. Delay to peak 1 was determined as the period between EOD onset and timing of peak 1. In addition, EOD frequency content was calculated by fast Fourier transformation.

We grouped EODs of *C. numenius* into three arbitrary types based on variation in duration (Long, Intermediate, and Short EOD), following a previous study that revealed extensive individual variation in EOD duration within this species (Paul et al., 2015). In the following analysis, we used the average value without separating types.

### Simultaneous recording and analysis of EOD and EOD command

This recording session was performed after EOD recording and before evoked potential recording only for three fish. Fish were anesthetized in a solution of 300 mg/L tricaine methanesulfonate (MS-222) (Sigma-Aldrich) and positioned on a plastic platform in a recording chamber filled with fresh water (Fig. 2B). Fish were restrained by lateral plastic pins and a plastic tube on the tail (Fig. 2B). Freshwater was provided through a pipette tip in the fish’s mouth. EOD commands from spinal EMNs were recorded with a pair of electrodes located within the plastic tube and oriented parallel to the fish’s EOD, amplified 1,000 times, and band-pass filtered (10 Hz–5 kHz) (Model 1700, A-M systems). While EOD commands from EMNs were recorded, the EODs were recorded by separate electrodes, amplified 10 times and band-pass filtered (1 Hz–50 kHz) (BMA-200, Ardmore). These recordings were digitized at a rate of 1 MHz and saved (TDS 3014C, Tektronix). We recorded 9–11 trials from each fish.

**Figure 2.**
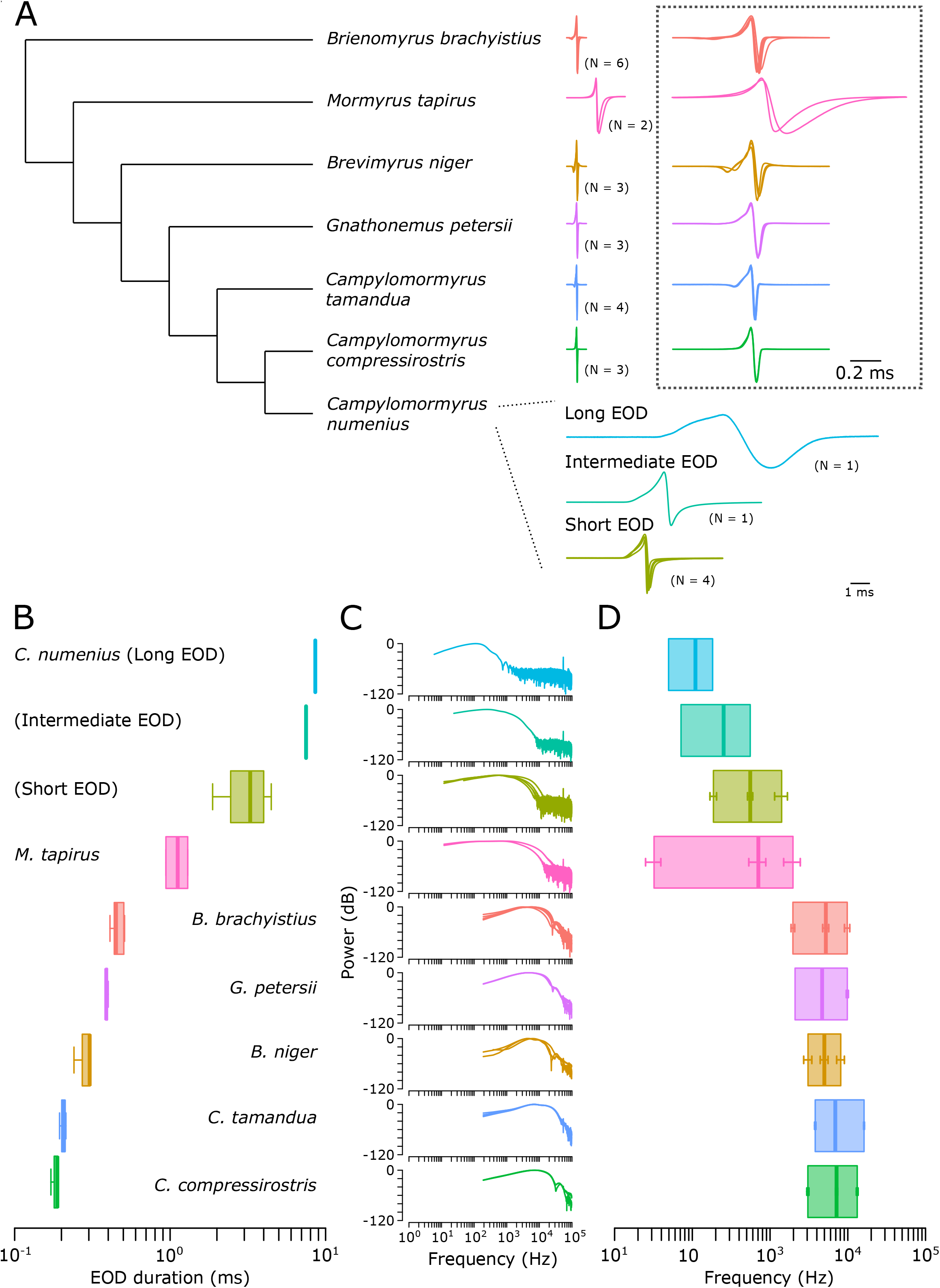
Mormyrids generate species-specific electric organ discharges (EODs). (A) Cladogram based on consensus trees of the species studied (Sullivan et al., 2000; Lavoué et al., 2003; Feulner et al., 2008; Sukhum et al., 2018). EOD traces are plotted as overlays of waveforms recorded from N individuals of each species and aligned to peak 1, defined as the head-positive peak. The EODs in *C. numenius* are displayed in three categories with distinct EOD durations (Long, Intermediate and Short). Expanded EODs for all other species are shown in the dotted box to the right. (B) Box plots of EOD durations from each species, sorted by EOD duration. (C) Power spectra of EODs from each species, from the same individuals shown in A. Each trace indicates the average EOD power spectrum from one individual. (D) Summary of EOD power spectra. Each bold line inside the box indicates the mean peak power frequency. The box limits indicate the mean lower and higher frequencies 3 dB below the peak. Error bars indicate the standard error of the mean.

EOD command traces from EMNs were averaged across trials, and EOD traces were filtered by a 21st-order median filter whose time window was 0.02 ms and averaged across trials. EOD onset was determined in the same way we determined EOD onset in freely swimming EOD recordings. Delay to EOD onset (D_Onset_) was calculated as the period between EOD command onset and EOD onset. Delay to peak 1 of EOD (D_P1_) from EOD command was calculated as the sum of D_Onset_ and the delay between EOD onset and peak 1 recorded from freely swimming fish.

### Surgery and evoked potential recording

This recording session was performed after EOD recording (or simultaneous recording of EODs and EOD commands for the three fish). We prepared fish for *in vivo* recordings from ELa and ELp as described previously (Carlson, 2009, Lyons-Warren et al., 2013a). Briefly, fish were anesthetized in 300 mg/L MS-222 and paralyzed with an intramuscular injection of 0.1 mL of 3.0 mg/mL gallamine triethiodide (Flaxedil) (Sigma-Aldrich). The fish were then transferred to a recording chamber (20 × 12.5 × 45 cm) filled with water and positioned on a plastic platform, leaving a small region of the head above water level. During surgery, we maintained general anesthesia by respirating the fish with an aerated solution of 100 mg/mL MS-222 through a pipette tip placed in the mouth. For *Campylomormyrus* species, we connected a hose made from heat shrink tubing to the pipette tip so as to provide respiration to the long, tube-like mouth. For local anesthesia, we applied 0.4% Lidocaine on the skin overlying the incision site, and then made an incision to uncover the skull overlying the ELa and ELp. Next, we glued a head post to the skull before using a dental drill and forceps to remove a rectangular piece of skull covering the ELa and ELp. In *Campylomormyrus* species, the ELa and ELp are not exposed superficially, so we exposed ELa and ELp by separating the optic tectum and the valvula cerebellum using two retractors made from borosilicate capillary glass (Vélez and Carlson, 2016; Vélez et al., 2017). After exposing ELa and ELp, we placed a reference electrode on the nearby cerebellum. Following surgery, we switched respiration to fresh water and allowed the fish to recover from general anesthesia. We monitored the anesthetized state of the fish with a pair of electrodes oriented parallel to its EO within a plastic tube to record fictive EOD commands produced by the EMNs (Carlson, 2009; Lyons-Warren et al., 2013a). These EOD commands were 1,000x amplified (Model 1700, A-M systems) and sent to a window discriminator for time stamping (SYS-121, World Precision Instruments). At the end of the recording session, the respiration of the fish was switched back to 100 mg/L MS-222 until no fictive EOD could be recorded, and then the fish was sacrificed by freezing.

Evoked potentials in ELa and ELp were obtained with glass microelectrodes made of borosilicate capillary glass (o.d. = 1.0 mm, i.d. = 0.5 mm; Model 626000, A-M Systems) pulled on a micropipette puller (Model P-97, Sutter Instrument Company), broken to a tip diameter of 10–20 μm and filled with 3M NaCl solution. Evoked potentials were 1,000x amplified, band-pass filtered (10 Hz–5 kHz) (Model 1700, A-M systems), digitized at a rate of 97.7 kHz (RX 8, Tucker Davis Technologies) and saved using custom software in Matlab (Mathworks).

### Sensory stimulation

We used three vertical electrodes on each side of the recording chamber (anodal to the fish’s left, cathodal to the right) to deliver transverse stimuli with normal polarity (peak preceding trough). Digital stimuli were generated using custom software in Matlab, converted to analog signals with a signal processor (RX8, Tucker Davis Technologies), attenuated with an attenuator (PA5, Tucker Davis Technologies) and isolated from ground with a stimulus isolation unit (Model 2200, A-M Systems).

To examine corollary discharge inhibition of sensory responses, we delivered 0.2 ms bipolar square pulses at several delays following the EOD command onset. The onset was determined as the first negative peak of the EOD command waveform that consists of a three-spike potential resulting from the synchronous activation of EMNs. Each delay was repeated 10 times and the averaged response was used for analysis. First, we recorded sensory responses in the ELp at delays between 0 and 20 ms in 0.5-ms steps. Based on these initial data, we used custom software written in R to determine the range of delays to examine corollary discharge inhibition with higher time resolution. The algorithm included the following steps: (1) calculated peak-to-peak amplitudes as a measure of response across all delays, (2) normalized all responses to the maximum response, (3) determined the latency resulting in the minimum response amplitude, (4) determined the onset and offset delays that resulted in responses just less than 80% of the maximum response, (5) determined the range of delays to examine corollary discharge inhibition with higher time resolution as 2 ms before the onset time to 2 ms after the offset time. Then, we recorded sensory responses in the ELp to stimulus delays across this range in 51 equally spaced steps (~ 0.1–0.2 ms). After recording in the ELp, we recorded sensory responses in the ELa using the same stimulus delays used in both the broad and narrow ranges tested in ELp. The stimulus sequences were randomized in all recordings.

To observe clear corollary discharge inhibition, we needed to decide on an adequate stimulus intensity for each tested individual before measuring of corollary discharge. First, we recorded evoked potentials at 0, 3, 4 and 5 ms delays at 20 dB attenuation (reference intensity of 736 mV/cm) and determined peak-to-peak amplitudes of each delay. Then, we calculated ratios of peak-to-peak amplitudes at 3, 4 and 5 ms delay to one at 0 ms delay. If the minimum ratio was under 30%, we chose this stimulus intensity. If the ratio was above 30%, we reduced the intensity by adding 5 dB attenuation and performed above recording until the ratio got under 30%. From this procedure, we chose 23.4 mV/cm for 1 *C. compressirostris* and 73.6 mV/cm for all the other fishes.

### Evoked potential recording analysis

We characterized corollary discharge inhibition with respect to timing and duration. All analyses here were done using the recordings from the high resolution, narrow range of stimulus delays. Normalized amplitude was calculated by the following steps: (1) calculated peak-to-peak amplitude for each delay, (2) subtracted by the minimum peak-to-peak amplitude across all delays and (3) divided by the maximum peak-to-peak amplitude across all delays, which leads to setting the maximum value as 1 and the minimum value as 0. Then, we set an 80% threshold to determine the inhibition onset, offset, and duration. In addition, we determined the peak time of inhibition as the stimulus delay at which the response amplitude was minimal.

### KO recording and analysis

KO recording data from *B. brachyistius* (n = 6 KOs), *B. niger* (n = 5), *Campylomormyrus alces* (n = 1), *C. compressirostris* (n = 7), *C. numenius* (n = 2) *and C. tamandua* (n = 2) came from previously published studies (Trzcinski, 2008; Trzcinski and Hopkins, 2008; Lyons-Warren et al., 2012; Baker et al., 2015). Based on a recent study of *Campylomormyrus* species (Paul et al., 2015), we concluded that *Campylomormyrus sp*. B shown in Trzcinski (2008) and Trzcinski and Hopkins (2008) was *C. numenius*.

The recording methods were generally shared among these previous studies. Similar to our evoked potential recording, fish were immobilized with Flaxedil, transferred to a recording chamber filled with freshwater and positioned on a plastic platform with lateral support. The fish were provided freshwater through a pipette tip in the fish’s mouth while monitoring the fish’s EOD command signals using a pair of electrodes placed next to the fish’s tail. Extracellular recordings of KO spikes were made using a wire electrode inside glass capillary tubing that was placed directly next to a KO. The signals were amplified, digitized, and recorded with custom software in Matlab. Sensory stimuli consisted of conspecific EOD waveforms generated in Matlab, digital-to-analog converted, attenuated, and delivered as a constant-current stimulus through the electrode.

These previous studies recorded KO responses to multiple waveforms at several intensities. For each recording, relative but not absolute intensities were known, because the amount of current going into the electroreceptor pore is dependent upon the position of the electrode tip relative to the pore, the size and shape of the pore, and the resistive paths between the electrode tip and the pore. Importantly, however, self-generated EODs will be of relatively large intensity. We therefore chose among these data here using the following criteria: (1) the EOD waveform stimulus was the inverted form of a conspecific EOD recorded with recording electrode at the head and reference electrode at the tail to simulate the self-generated EOD waveform; and (2) the largest stimulus intensity tested that did not result in a stimulus artifact that exceeded KO spike amplitude.

To calculate normalized KO responses, we made peristimulus time histograms (bin size = 0.02048 ms) of KO spikes and normalized them by the maximum spike counts. Delay to peak KO response was determined as the period between the onset of the EOD stimulus and the time of the maximum KO response.

### Experimental design and statistical analysis

The primary objective of this study was to determine the relationship between EOD waveform and corollary discharge inhibition. We used 7 species including 27 fishes, and recorded EODs from freely swimming individuals followed by evoked potential recording from the midbrain. Subsequently, we asked whether corollary discharge inhibition overlapped KO spike timing to block responses to self-generated EODs. To compare the time courses between corollary discharge inhibition and EOD using an identical reference of EOD command onset, we performed simultaneous recording of EOD and EOD command from a subset (3 species including 3 fishes) of them before performing electrophysiology. Furthermore, we measured KO response latency to self-generated EODs using previously published data (Trzcinski, 2008; Trzcinski and Hopkins, 2008; Lyons-Warren et al., 2012; Baker et al., 2015).

For statistical analysis, we used a phylogenetic generalized least squares (PGLS) model with a Brownian correlation structure to account for phylogenetic effects on the correlation analyses. We used a previously constructed bootstrapped maximum-likelihood tree from 73 Cytb osteoglossomorph sequences (Sukhum et al., 2018). To include data from species that have not been sequenced, we used sequence data from within monophyletic genera and chose the species sequence with the shortest phylogenetic distance from the genus node. In this analysis, we used average values within species of EOD waveform, corollary discharge inhibition and KO response. Note that regression line, t value, and p value were calculated with this PGLS model whereas correlation coefficients were calculated with Pearson’s method. All phylogenetic analyses were performed in R programming software with the ape and nlme packages (Paradis and Schliep, 2018; Pinheiro et al., 2020).

## RESULTS

### Mormyrids have diverse species-specific EODs

To relate corollary discharge inhibition to EOD waveform, we recorded EODs individually from 7 species before performing evoked potential recordings (Fig. 2A). EOD duration varied widely across species, from as short as 0.17 ms in *C. compressirostris* to as long as 8.59 ms in *C. numenius* (Fig. 2B). Fast Fourier transformation revealed that peak power frequencies ranged from 110 Hz in *C. numenius* to 7610 Hz in *C. compressirostris* (Fig. 2C, D).

### Corollary-discharge timing and duration varies among species

To measure the inhibitory effect of corollary discharge in the KO pathway, we performed evoked potential recordings from ELa and ELp (Fig. 3A). We stimulated with 0.2-ms bipolar square electric pulses delivered with a delay of 0–20 ms following the EOD command from spinal electromotor neurons (EMNs) (Fig. 3A). To our knowledge, these are the first evoked potential recordings from the midbrain of *Campylomormyrus* species and *M. tapirus*. The recording traces of evoked potentials from those species were similar to those of previously reported species, including *B. brachyistius, B. niger, G. petersii, Petrocephalus microphthalmus* and *Petrocephalus tenuicauda* (Russell and Bell, 1978; Amagai, 1998; Carlson, 2009; Lyons-Warren et al., 2013b; Vélez and Carlson, 2016): electrosensory stimulation elicited sharp and short-latency (~2–4 ms) evoked potentials in ELa, and broad and longer-latency (~6–10 ms) evoked potentials in ELp (Fig. 3B).

**Figure 3.**
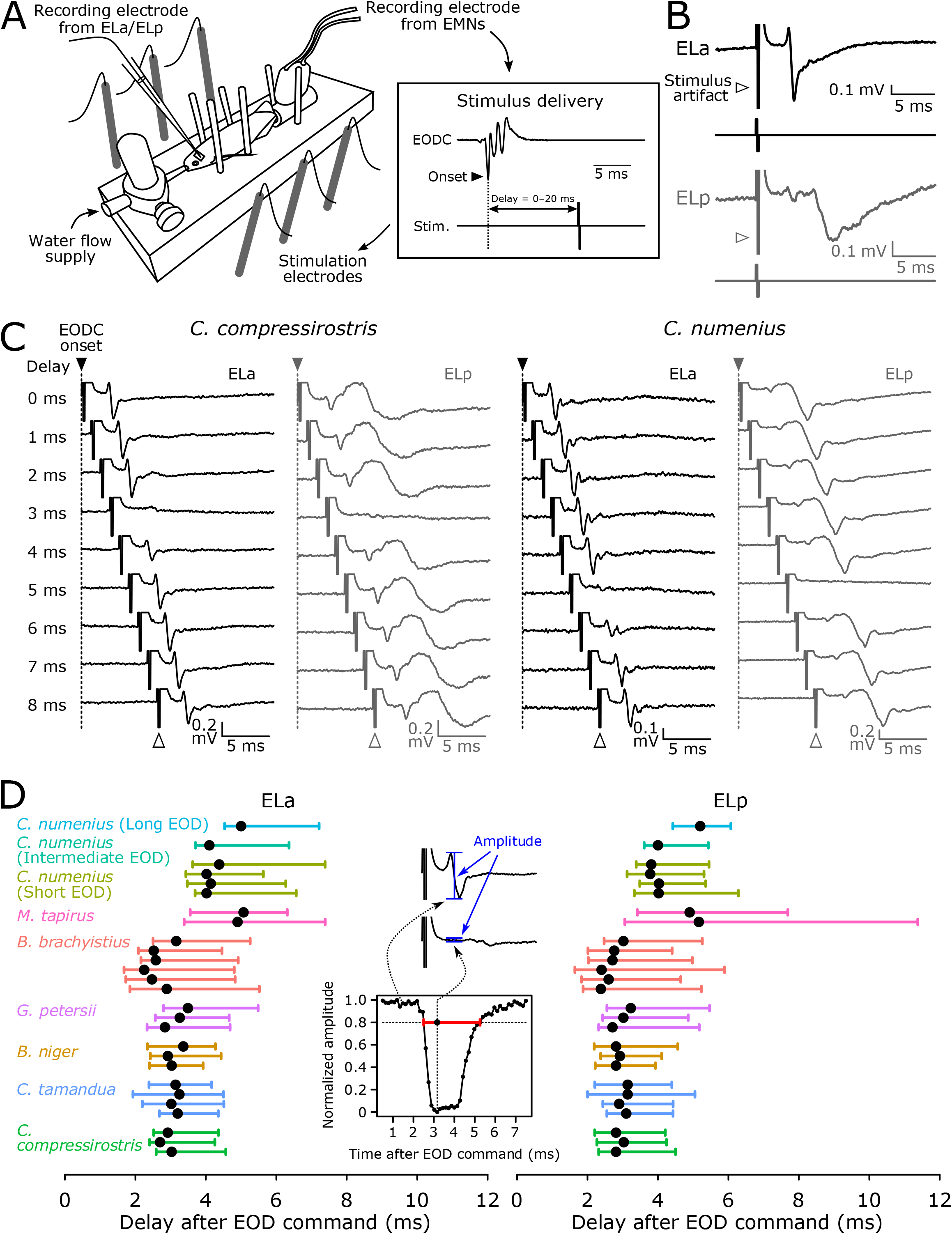
Corollary discharge inhibition in the communication circuit varies in timing and duration among mormyrid species. (A) Schematic representation of electrophysiological recording from the fish. An extracellular electrode inside a tube placed over the tail records EOD commands from spinal electromotor neurons (EMN). To assess corollary discharge inhibition related to EOD production, we delivered sensory stimuli at different delays (0–20 ms) after the EOD command (EODC) onset, which is determined as the first negative peak (indicated by black arrowhead), while recording evoked potentials in ELa or ELp. (B) Representative mean evoked potentials (n = 10 traces) obtained from ELa and ELp in *M. tapirus*. (D) Representative mean evoked potentials in response to stimuli at varying delays following the EOD command (0–8 ms) in *C. compressirostris* and *C. numenius* (Long EOD type). (E) Summary of corollary discharge inhibition across species. Inset describes the measurement of corollary discharge inhibition timing. The response magnitudes of evoked potentials were calculated as the peak-to-peak amplitude (blue bars) and normalized to the maximum peak-to-peak amplitude across all stimulus delays. Using an 80% threshold, we determined inhibition onset, offset, and duration (red bar). The large point on the red bar indicates the peak inhibition time. Left and right panels show the inhibition periods relative to the EOD command in ELa and ELp, respectively, across all individuals studied.

In each species, we found a narrow range of stimulus delays for which electrosensory responses were blocked by corollary discharge inhibition (Fig. 3C), as shown previously in *B. brachyistius* and *B. niger* (Amagai, 1998; Lyons-Warren et al., 2013b; Vélez and Carlson, 2016). From these evoked potential traces, we determined the corollary discharge inhibition window for each individual using normalized amplitudes and an 80% cutoff line (Fig. 3D). This revealed that corollary discharge onset, offset, duration, and peak time varied among species (Fig. 3D). For example, evoked potentials in the short-EOD *C. compressirostris* were blocked when sensory stimuli were delivered with a 3-4 ms delay following the EOD command, whereas evoked potentials in the long-EOD *C. numenius* were blocked when stimuli were delivered with a 5-6 ms delay following the EOD command (Fig. 3C).

### Corollary discharge timing is correlated with EOD waveform

We next asked whether species diversity in EOD waveform (Fig. 2) was correlated with species diversity in corollary discharge timing (Fig. 3). First, we tested the relationship between EOD duration and corollary discharge inhibition duration (Fig. 4A, B). Inhibition duration in ELa was positively correlated with EOD duration (Fig. 4A; Pearson’s correlation coefficient, 0.4164; slope, 0.1957; intercept, 2.4221; t = 5.1390; *p* = 0.0036), whereas inhibition duration in ELp was not correlated with EOD duration (Fig. 4B; Pearson’s correlation coefficient, −0.0536; slope, 0.0061; intercept, 3.3004; t = 0.0534; *p* = 0.9595).

**Figure 4.**
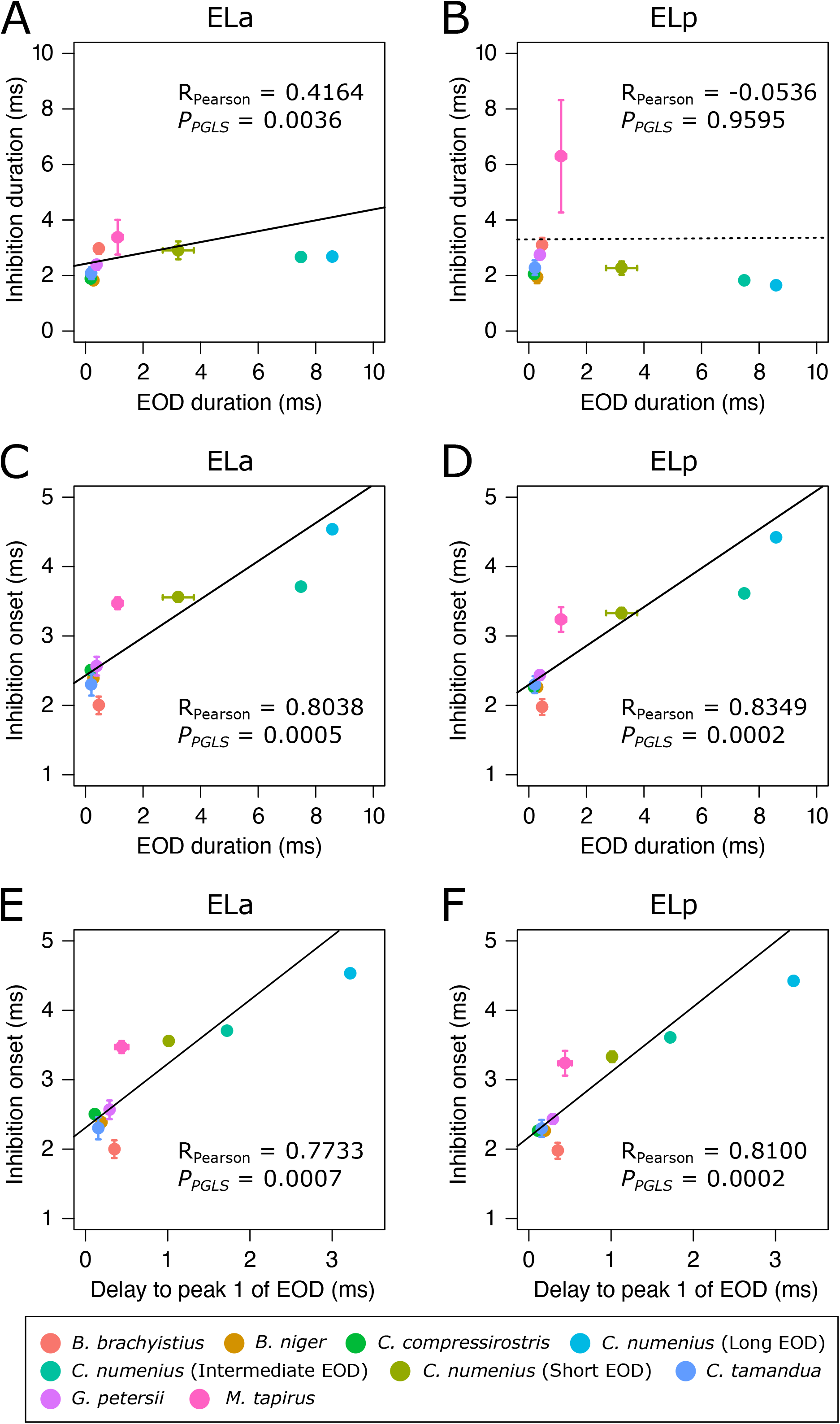
Corollary discharge inhibition onset is correlated with EOD duration. (A, B) Plots of inhibition duration (y axis) against EOD duration (x axis) in ELa and ELp. (C, D) Plots of inhibition onset against EOD duration in ELa and ELp. (E, F) Plots of inhibition onset against delay to peak 1 of the EOD in ELa and ELp. Points show the mean value of species ± the standard error of the mean (but not for *C. numenius*). Regression lines were determined using a PGLS analysis. Significant correlations are indicated by a solid line and insignificant correlations are indicated by a dotted line. Although *C. numenius* is represented as three groups (long, intermediate and short EOD), the regression was calculated using average values of each species. *R_Pearson_*, Pearson’s correlation coefficient; *P_PGLS_*, *p*-value calculated from PGLS.

The different timing of corollary discharge inhibition in species with different EOD durations (Fig. 3C, D) suggested that corollary discharge onset rather than duration might be associated with EOD duration. Indeed, we found that inhibition onset was strongly correlated with EOD duration in both ELa (Fig. 4C; Pearson’s correlation coefficient, 0.8162; slope, 0.2747; intercept, 2.4306; t = 7.9658; *p* = 0.0004) and ELp (Fig. 4D; Pearson’s correlation coefficient, 0.8451; slope, 0.2798; intercept, 2.2976; t = 10.0555; *p* = 0.0001).

Here we recall two important features of KO responses to self-generated EODs: (1) each receptor responds with time-locked spikes to outside-positive changes in voltage across the skin; and (2) all KOs receive the same EOD waveform, which is inverted in polarity compared to the head-positive EOD recordings shown in Fig. 2. This suggests that KOs should respond immediately after peak 1 of the EOD, when the self-generated EOD transitions from a negative to positive peak. This further suggests that, in order to effectively block responses to self-generated EODs, the timing of corollary discharge inhibition should also relate to the timing of EOD peak 1. Indeed, we found that inhibition onset was strongly correlated with the delay to EOD peak 1 in both ELa (Fig. 4E; Pearson’s correlation coefficient, 0.7733; slope, 0.9203; intercept, 2.3092; t = 7.4537; *p* = 0.0007) and ELp (Fig. 4F; Pearson’s correlation coefficient, 0.8100; slope, 0.9391; intercept, 2.1734; t = 9.3725; *p* = 0.0002).

Since our definition of EOD duration requires arbitrary cutoffs to define the beginning and end of the EOD, we also determined whether EOD peak power frequency correlated with inhibition duration and onset. We found a significant negative correlation between peak power frequency and inhibition duration in ELa (Pearson’s correlation coefficient, −0.7585; slope, −0.0001; intercept, 3.1375; t = −7.1453; *p* = 0.0008), but this relationship was not significant in ELp (Pearson’s correlation coefficient, −0.5430; slope, −3.54 × 10^−5^; intercept, 3.4525; t = −0.4669; *p* = 0.6602). By contrast, there were significant negative correlations between peak power frequency and inhibition onset in both ELa (Pearson’s correlation coefficient, −0.8875; slope, −0.0002; intercept, 3.4186; t = −11.5879; *p* = 0.0001) and ELp (Pearson’s correlation coefficient, −0.9026; slope, −0.0002; intercept, 3.2987; t = −14.8227; *p* = 2.53 × 10^−5^).

### KO responses to self-generated EODs depend on the delay to peak 1

We tested the hypothesis that KO responses are time-locked to peak 1 of the EOD in both short-EOD and long-EOD species. We used previously published data from 6 species (Trzcinski, 2008; Trzcinski and Hopkins, 2008; Lyons-Warren et al., 2012; Baker et al., 2015) and examined the timing of KO spiking responses to conspecific EODs. By convention, ‘normal’ EOD polarity is defined as a waveform recorded with the recording electrode at the head and a reference electrode at the tail as shown in Fig. 2. Here we focused on KO responses to ‘inverted’ EODs that represent the same waveform that KOs receive in response to self-generated EODs. We found that KOs responded with time-locked spikes with short delay following peak 1 of the EOD (Fig. 5A; delays between peak 1 of EOD and peak KO response were 0.14 ms in *C. compressirostris*, 0.07 ms in *B. brachyistius* and 0.37 ms in *C. numenius*). Across species, delay to peak KO response strongly correlated with delay to peak 1 of the EOD (Fig. 5B; Pearson’s correlation coefficient, 0.9937; slope, 1.1550; intercept, 0.1128; t = 41.1242; *p* = 2.10 × 10^−6^).

**Figure 5.**
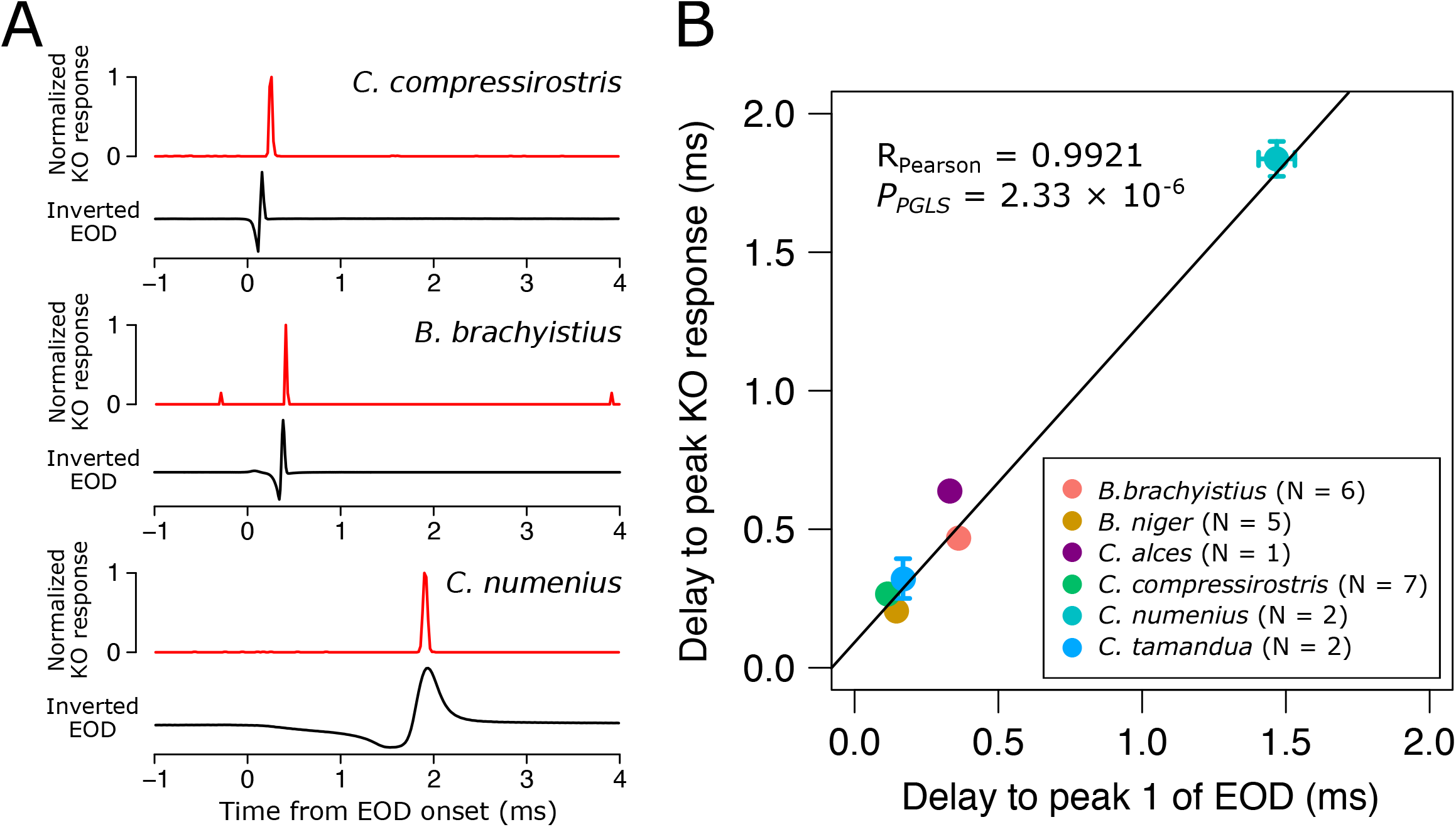
Knollenorgans respond with time-locked spikes following peak 1 of EOD stimuli that simulate self-generated EODs. (A) Example traces of normalized KO responses of *C. compressirostris, B. brachyistius*, and *C. numenius*. Lower traces show the inverted EODs of conspecifics whose onsets are aligned to time 0. (B) Plots of delay to peak KO response against delay to peak 1 of EOD. Points show the mean value of species ± the standard error of the mean. Regression line was determined using a PGLS analysis. R_Pearson_, Pearson’s correlation coefficient; *P_PGLS_*, *p*-value calculated from PGLS.

### Corollary discharge inhibition timing blocks KO responses to self-generated EODs

Our results so far revealed that corollary discharge inhibition and KO spiking responses were both correlated with delay to peak 1 of the EOD. This leads to the further question of whether the time-shifted corollary discharge inhibition actually blocks response to self-generated EODs. To address this question, we measured the delay between the EOD command from spinal motor neurons and the EOD in fish that were not electrically silenced and paralyzed. We found that delays between EOD command onset and EOD onset were similar among the three species (*C. compressirostris*, 3.12 ms; *B. brachyistius*, 3.08 ms; *C. numenius*, 3.28 ms) (Fig. 6). Thus, the delays between EOD command onset and peak 1 of the EOD varied among the species (*C. compressirostris*, 3.24 ms; *B. brachyistius*, 3.41 ms; *C. numenius*, 5.05 ms) (Fig. 6). Comparing the time courses of the EOD and corollary discharge inhibition using an identical reference of the EOD command onset revealed that corollary discharge inhibition overlapped with the timing of EOD peak 1 across species (Fig. 6).

**Figure 6.**
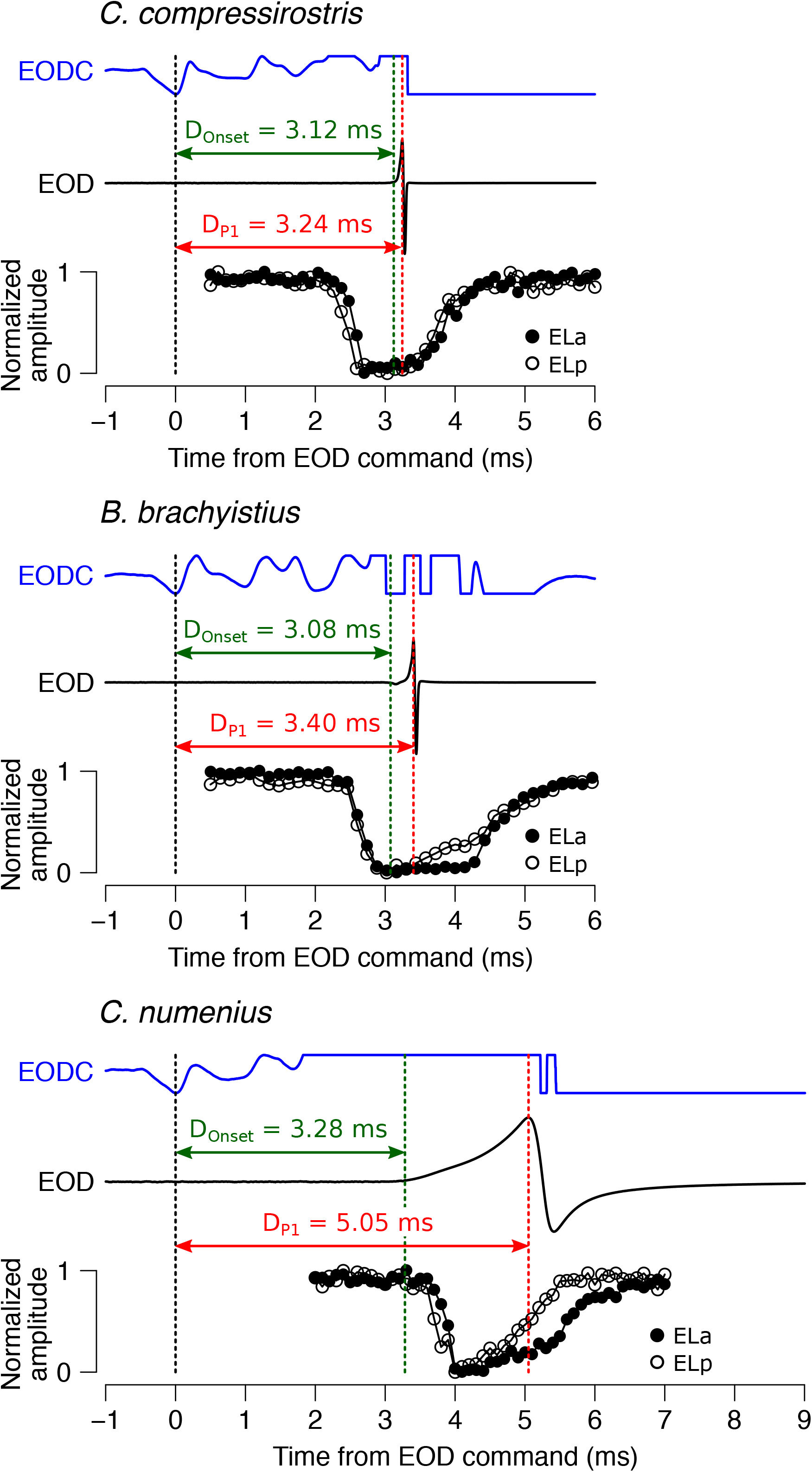
Corollary discharge inhibition is timed to block responses to peak 1 of the EOD. We compared the time courses of the EOD command (EODC) (upper blue traces), the EOD (middle traces), and corollary discharge inhibition. Vertical dotted lines indicate EODC onset (black), EOD onset (green), and peak 1 of the EOD (red). D_onset_, Delay to EOD onset; D_P1_, Delay to peak 1 of EOD.

### Corollary discharge inhibition timing is correlated with individual EOD waveform variation within *C. numenius*

The high degree of individual variation in EOD duration in *C. numenius* (Fig. 2A, B) facilitates an examination of the correlation between EOD waveform and corollary discharge within species. Similar to our results across species, corollary discharge onset was strongly correlated with delay to peak 1 of the EOD in both ELa (Fig. 7A; Pearson’s correlation coefficient, 0.9548, slope = 0.4307, intercept = 3.0983, t = 6.4206, *p* = 0.0030) and ELp (Fig. 7B; Pearson’s correlation coefficient, 0.9460, slope = 0.4802, intercept = 2.8388, t = 5.8355, *p* = 0.0043).

**Figure 7.**
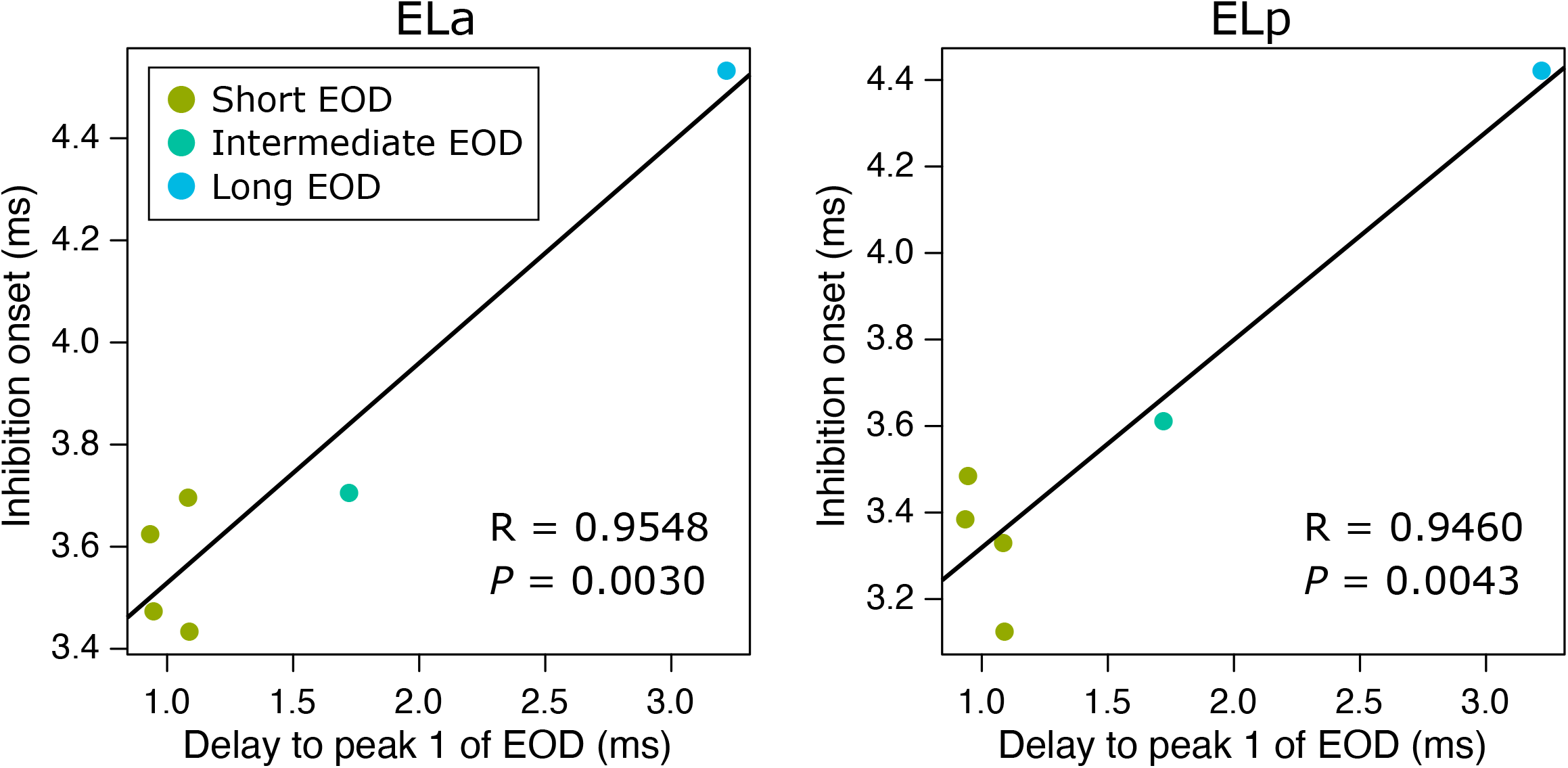
Corollary discharge onset is correlated with individual EOD waveform variation among *C. numenius*. Plots of inhibition onset (y axis) against delay to peak 1 of EOD (x axis) in ELa and ELp. Points show individual values. Regression lines were determined using a linear regression analysis. R, Pearson’s correlation coefficient; *P*, *p*-value of the correlation.

## DISCUSSION

Our findings provide evidence that diverse communication signals in mormyrids are correlated with corollary discharge inhibition of the electrosensory pathway. We show that EOD duration is only weakly correlated with the duration of corollary discharge inhibition, but strongly correlated with the onset of corollary discharge inhibition (Fig. 4). Taking into account that electroreceptors (KOs) produce spikes with short latency following peak 1 of the EOD (Fig. 5) and that the corollary discharge inhibition overlaps this peak (Fig. 6), we conclude that corollary discharge inhibition has evolved to shift its time window so as to optimally block KO spikes in response to self-generated EODs (Fig. 8).

**Figure 8.**
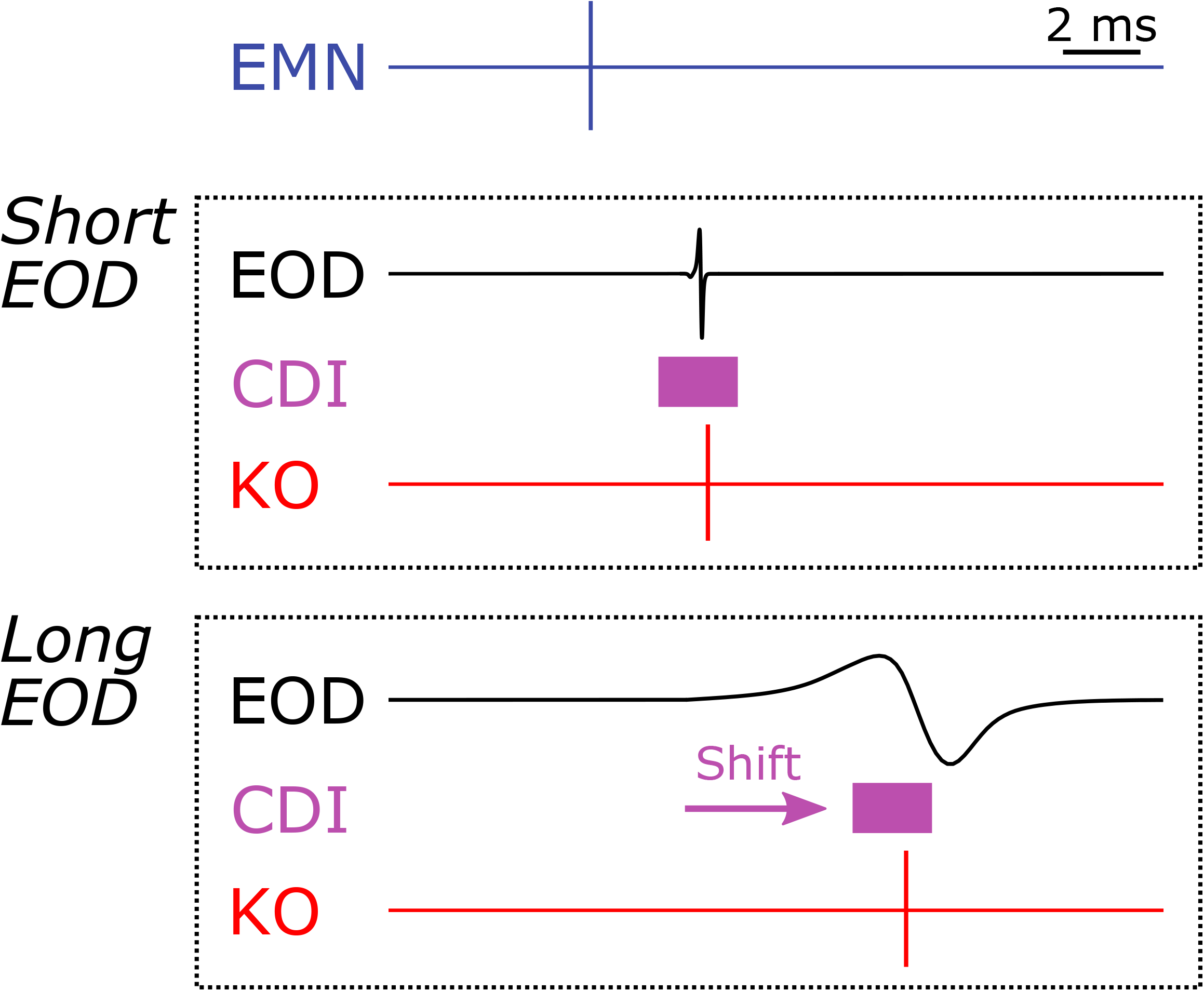
Time shift of corollary discharge inhibition underlies communication signal evolution in mormyrid. (A) Summary of corollary discharges between mormyrids with short-duration EOD and long-duration EOD. The schematic diagram shows spike timings of EMN and KO as well as EOD. Purple rectangle shows time window of corollary discharge inhibition (CDI).

Evolutionary change in behavior can result from changes to sensory systems, motor systems, or both (Katz, 2011, 2016; Martin, 2012; Stöckl and Kelber, 2019). Correlated evolution of sensory and motor systems is especially apparent in communication systems, as this requires evolutionary change in both signal production and reception (Bass and Hopkins, 1984; Bass, 1986; Otte, 1992; ter Hofstede et al., 2015; Silva and Antunes, 2017). Indeed, mormyrids show correlated evolution between senders and receivers of their electric communication signals: the frequency tuning of KOs is related to the frequency spectrum of conspecific EODs (Bass and Hopkins, 1984; Lyons-Warren et al., 2012; but see also Baker et al., 2015). Here we add a further insight that signal evolution accompanies evolutionary change of neural circuitry underlying sensorimotor integration. Corollary discharges that filter sensory responses to self-generated signals are ubiquitous across communicating animals (Crapse and Sommer, 2008). In addition, signals among related species often vary widely in temporal features (Otte, 1992; Hopkins, 1999; Podos, 2001), and changing the timing of communication signals alters the timing of reafferent input. Therefore, we expect similar evolutionary change in corollary discharge timing to be widespread across sensory modalities and taxa. For example, the temporal structures of species-specific songs of crickets are similarly diverse to mormyrid EODs (Otte, 1992; ter Hofstede et al., 2015), and a similar corollary discharge inhibition of the auditory pathway has been described in one species (Poulet and Hedwig, 2006).

Why does evolutionary change of EOD duration relate to the delay of corollary discharge inhibition rather than inhibition duration? Theoretically, it is possible to alter the duration to cover the entirety of EODs with different durations. However, changing the delay of corollary discharge inhibition without expanding the duration would avoid unnecessarily elongating the resulting insensitive period. This is because the KOs responding to self-generated EODs produce spikes only over a narrow window of time just after EOD peak 1 regardless of EOD duration (Fig. 5). Accordingly, we suggest an optimal evolutionary strategy for modifying corollary discharge inhibition during signal evolution: minimizing the inhibitory window to only what is necessary for blocking receptor responses.

Mormyrids have additional electrosensory systems, ampullary and mormyromast, used for passive and active electrolocation, respectively. Corollary discharge plays a significant, but different, role in these systems (von der Emde and Bell, 2003; Warren and Sawtell, 2016). The primary afferents of ampullary receptors are spontaneously active and exhibit long lasting spiking responses that generally peak more than 20 ms after the EOD (Bell and Russell, 1978). The primary afferents of mormyromasts have less spontaneous activity, but exhibit long lasting responses consisting of 2–8 spikes ~2 ms after the EOD (Bell, 1990b). Corollary discharge in both systems works to subtract predictive sensory consequences of reafference by activating a modifiable efference copy (i.e. ‘negative image’) through cerebellar-like circuitry in the ELL cortex (Warren and Sawtell, 2016).In this circuit, the negative image is generated in synapses between granule cells forming parallel fibers and principal cells through spike-timing-dependent plasticity (Bell et al., 1997). While the granule cells receive stereotyped corollary discharge inputs with short delays after the EOD command, their outputs are more temporally diverse and delayed (Kennedy et al., 2014). This is an important feature to provide a temporal basis for generating a sufficiently long negative image (~200 ms). Species with longer EOD durations likely have longer-lasting responses, which would require even more temporal dispersion among granule cells to generate a longer-lasting negative image. In addition, through a separate pathway, corollary discharge input facilitates responses to afferent input from mormyromasts, thereby selectively enhancing responses to reafferent EODs (Bell, 1990a). Here, too, species and individual differences in EOD duration may require corollary discharge input with different time courses.

What might be the source of species differences in the delay of corollary discharge inhibition of the KO pathway? The command nucleus (CN) drives the EO to produce each EOD through the medullary relay nucleus (MRN) and the EMN (Fig. 1A). The CN also provides corollary discharge inhibition to the nELL through the bulbar command-associated nucleus (BCA), the mesencephalic command-associated nucleus (MCA), and the sublemniscal nucleus (slem) (Fig. 1A). Note that the EOD command waveform recorded from the EMN is almost identical across species and independent of EOD duration (Fig. 5; Bass and Hopkins, 1983), suggesting that command circuitry does not contribute to corollary discharge delay. Thus, the corollary discharge pathway must be adjusting the corollary discharge delay (Fig. 1A; Bell et al., 1983). The BCA and MCA are interesting candidates because they are involved in controlling the inter-pulse interval (IPI) between EODs and longer EODs require longer IPIs (von der Emde et al., 2000; Carlson, 2002a, 2002b, 2003). Future studies will compare this corollary discharge pathway across species to identify the source of species differences in delay.

What might be the mechanism for species differences in the corollary discharge inhibition delay? There are at least three types of modifications that could change this delay. First, elongating axons could increase transmission delays much like the ‘delay lines’ observed in several sensory systems (Carr and Konishi 1988; Lyons-Warren et al., 2013b). Second, decreasing myelination and/or smaller axonal diameter could reduce the conduction velocity of axonal action potential propagation (Waxman et al., 1972; Seidl, 2014). Third, intrinsic properties of neurons (e.g. A-current, kinetics of transient outward K^+^ current) can also affect inhibition timing (Getting, 1983). A combination of physiological recording and anatomical characterization of the corollary discharge pathway across species will uncover the mechanistic basis for evolutionary change in corollary discharge inhibition delays.

In addition to species differences, we show that individual differences in corollary discharge delay are correlated with delay to EOD peak 1 in *C. numenius* (Fig. 6). A previous study demonstrated that EOD duration changes substantially with growth in *C. numenius*, and this correlates with ontogenetic changes in EO anatomy (Paul et al., 2015). For many mormyrid species, EOD waveform varies with ontogeny, sex, relative dominance, and season (Bass, 1986; Hopkins, 1999; Carlson et al., 2000; Werneyer and Kramer, 2006). The existence of individual variation in corollary discharge delays raises the question whether the same mechanism is used for individual variation of corollary discharge as for species differences. Moreover, are changes in corollary discharge timing and EOD duration mediated by a shared central regulatory pathway, or through neuronal plasticity associated with changing sensory input in response to self-generated EODs?

There are at least three possibilities that might explain how corollary discharge delay changes along with individual or evolutionary change in EOD waveform: (1) EOD command pathway drives both changes in EOD waveform and corollary discharge delay; (2) genetic regulation drives both changes; (3) neural plasticity adapts the corollary discharge to changes in EOD duration. (1) would be impossible because there is no way for the electromotor network to provide the corollary discharge pathway with waveform information. EOD waveform is determined by the biophysical characteristics of electrocytes (Bennett, 1971; Bass, 1986), independent from the EOD command (Fig. 5; Bass and Hopkins, 1983). (2) would be possible. Recently, the genomic basis of EO anatomy and physiology is being increasingly well studied (Gallant et al., 2014; Gallant and O’Connell, 2020). It would be interesting if a central regulatory mechanism lead to correlated transcriptional changes in the EO and corollary discharge pathway. (3) would be possible. Although previous studies suggest that corollary discharge in the KO system is invariant over several hours of electrophysiological experimentation, this was under a limited, unnatural situation in which the EOD was absent and no association with an alternative EOD was tested (Bell and Grant, 1989). Such an associative mechanism possibly takes place in the nELL as it is only site at which corollary discharge and KO sensory processing converge (Mugnaini and Maler, 1987). However, it is possible that retrograde signals could make an association at earlier stages in the corollary discharge pathway.

Here we show evolutionary change of corollary discharge inhibition for the first time. Our results strongly suggest a corollary discharge pathway makes an appropriate delay to block receptor responses to self-generated signals. Future studies will seek to identify the source of delays and the cellular mechanisms using electrophysiological and anatomical approaches. Furthermore, it will be interesting to study time shifts of corollary discharge inhibition during signal development within individuals. Such studies will reveal the mechanisms by which sensorimotor integration is adjusted to account for species and individual differences in behavior.

## Acknowledgement

This work was supported by the National Science Foundation (Grant IOS-1755071 to B.A.C.) and the Uehara Memorial Foundation (to M.F.). We thank Carl D. Hopkins and Natalie Trzcinski for kindly providing their Knollenorgan recording data; Adalee Lube for assistance with electrophysiology; and Erika Schumacher for assistance with statistical analysis.

## Author Contributions

M.F. and B.A.C. designed research; M.F. performed research; M.F. analyzed data; M.F. and B.A.C. wrote the paper.

## Notes

**Conflict of Interest Statement:** None

### Competing Interest Statement

The authors have declared no competing interest.

